# Unveiling common transcriptomic features between melanoma brain metastases and neurodegenerative diseases

**DOI:** 10.1101/2024.01.16.575868

**Authors:** Irene Soler-Sáez, Alcida Karz, Marta R. Hidalgo, Borja Gómez-Cabañes, Adolfo López-Cerdán, José F. Català-Senent, María de la Iglesia-Vayá, Eva Hernando, Francisco García-García

**Affiliations:** Computational Biomedicine Laboratory, Principe Felipe Research Center (CIPF), 46012, Valencia, Spain; Department of Pathology, New York University Grossman School of Medicine, New York, NY 10016, US; Biomedical Imaging Unit FISABIO-CIPF, Fundación para el Fomento de la Investigación Sanitaria y Biomédica de la Comunidad Valenciana, 46012, Valencia, Spain

## Abstract

Melanoma represents a critical clinical challenge due to its high incidence rates and unfavorable clinical outcomes. This type of skin cancer presents unique adaptability to the brain microenvironment, but its underlying molecular mechanisms are poorly understood. To further characterize its tumor neurobiology, we explore the relation between the transcriptional profiles of melanoma brain metastasis (MBM) and the neurodegenerative diseases Alzheimer’s disease, Parkinson’s disease, and multiple sclerosis. Through an *in silico* approach, we unveiled the neurodegenerative signature of MBM when compared to melanoma non-brain metastasis (53 dysregulated genes enriched in 11 functional terms) and to non tumor-bearing brain controls (195 dysregulated genes, mostly involved in development and cell differentiation, chromatin remodeling and nucleosome organization, and translation). Two genes, ITGA10 and DNAJC6, emerged as key potential markers, as they are dysregulated in both scenarios. Lastly, we developed a user-friendly web tool (https://bioinfo.cipf.es/metafun-mbm/) as an open source, so that any user can interactively delve into the results.

## INTRODUCTION

Brain metastasis is an urgent clinical problem in metastatic melanoma patients due to their incidence rates and poor outcomes. Melanoma is the third most common source of brain metastases, exceeded only by lung and breast adenocarcinomas (PMID: 27843591), and is the solid tumor with the highest propensity for homing to the brain (PMID: 33908612; PMID: 12173339). To successfully seed a secondary tumor, metastatic cells from extracranial tumors must pass through the ultra-selective blood-brain barrier. Then, adaptation to the brain microenvironment (BME) requires arbitration with unique cell types like astrocytes and microglia, which may be initially hostile and eventually supportive (PMID: 31601988; PMID: 33479058).

Melanoma’s relative adaptability to the challenges of the BME is thought to stem from melanoma cells’ developmental origin from the neural crest. In fact, analysis of single cell gene expression data from treatment-naive tumors shows that melanoma brain metastatic cells adopt neural-like signatures (PMID: 35803246). Additional evidence points to a link between brain metastasis and neurodegenerative diseases. Melanoma cells have been shown to secrete the Alzheimer’s disease (AD) associated peptide amyloid B in order to suppress local inflammation in the BME (PMID: 35262173). Furthermore, a significant correlation between incidence of melanoma and Parkinson’s disease (PD) has been repeatedly observed (PMID: 32210791; PMID: 36788297), with both melanoma patients appearing to have an increased risk for PD and vice versa (PMID: 34215604). These two neurodegenerative conditions and MS, involve major inflammatory responses impacting brain resident cells, such as astrocytes and microglia, which have also been directly implicated in the adaptation of cancer cells to the brain parenchyma.

Despite these associations, an explicit, unbiased comparison between melanoma gene signatures and those of neurodegenerative diseases has not yet been conducted. We undertook this exercise to identify mechanisms undergone by melanoma cells as they adapt to the brain. In particular, we posit that efforts to evaluate the relation between melanoma brain metastasis (MBM) gene signatures and those of AD, PD and multiple sclerosis (MS) may expand our understanding of the biology of these tumors. Pathways dysregulated across CNS pathologies may be reflective of metabolic or immune adaptations to injury and thus represent potential vulnerabilities that could be targeted for therapeutic purposes. These pathways and the corresponding nominated targets should be assessed experimentally in follow-up studies to demonstrate their functional involvement in MBM.

## RESULTS

In order to explore the relationship between neurodegenerative gene signatures and those characteristic of melanoma brain metastasis (MBM), we followed an *in silico* approach. First, we identified published transcriptomic profiles of MBM, and brain expression datasets from patients with Alzheimer’s disease (AD), Parkinson’s disease (PD), and multiple sclerosis (MS). After performing differential expression analyses and meta-analyses of neurodegenerative diseases datasets, we identified a set of dysregulated genes shared with MBM, and the corresponding pathways and biological functions enriched. All the results obtained are available as an open resource on the web tool (https://bioinfo.cipf.es/metafun-mbm/). Our analysis supports the emerging notion that melanoma cells colonizing the brain adopt gene signatures reminiscent of neural cells altered by degenerative diseases (Figure 1).

**Figure 1.**
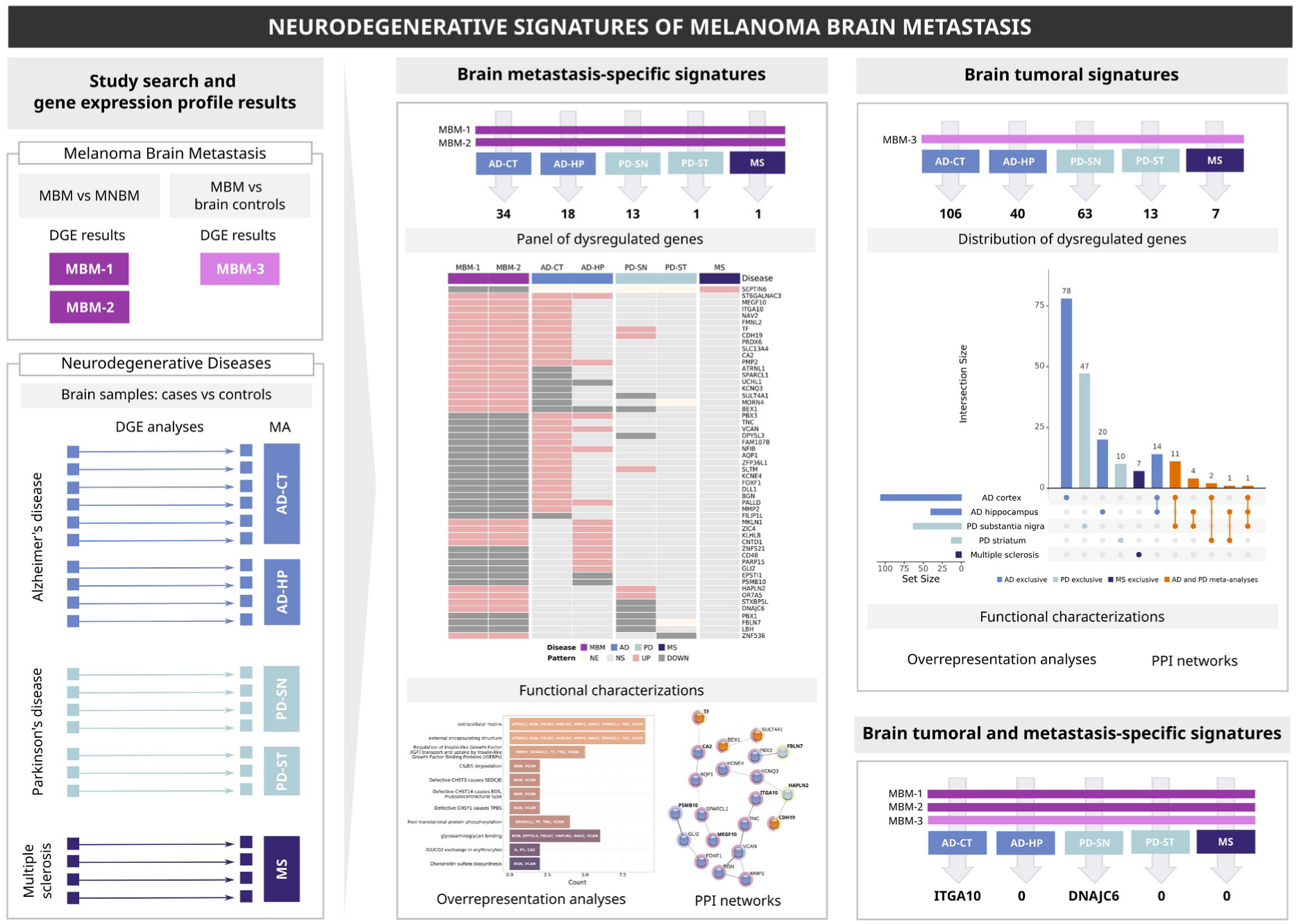
Implemented framework and overview of the work. Publicly available transcriptomic datasets were retrieved, and differential gene expression analyses downloaded between MBM vs MNBM or MBM vs non tumor-bearing brain controls. Similarly, for neurodegenerative diseases, transcriptomic datasets were downloaded, and differential gene expression analyses and meta-analyses were performed. A neurodegenerative gene signature characteristic of melanoma brain metastasis (i.e., MBM vs MNBM comparison) was identified and additional pathway analyses were conducted. A similar approach was implemented to draw aMBM tumor signature by comparing MBM vs control brains. Shared dysregulated genes across both scenarios were also unveiled. MBM: melanoma brain metastasis; MNBM: melanoma non-brain metastasis. AD: Alzheimer’s disease; CT: cortex; HP: hippocampus PD: Parkinson’s disease; SN: substantia nigra; ST: striatum; MS: multiple sclerosis; DGE: differential gene expression; MA: meta-analyses; NE: not evaluated; NS: not significant.

### Data collection and differential gene expression analyses by neurodegenerative disease

Literature screening led to the identification of public transcriptomic studies for the diseases of interest. We selected 3 melanoma patient datasets, hereby referred to as MBM-1, MBM-2, and MBM-3, and we retrieved the differential gene expression results. The first two studies identified differences in MBM versus non-brain melanoma metastases (MNBM), whileMBM-3 compared gene expression profiles of MBM vs healthy brain regions. Further details are presented in Table 1.

**Table 1.** Descriptive characteristics of the selected MBM studies. Upregulated genes refer to the number of significant genes with higher expression in the first term of the comparison (logFC > 0), and downregulated genes to those more highly expressed in the second term of the comparison (logFC < 0). MBM: melanoma brain metastasis; MNBM: melanoma non-brain metastasis; FFPE: Formalin-Fixed Paraffin-Embedded; single nuclei RNA-seq.

|  | **MBM-1** | **MBM-2** | **MBM-3** |
| --- | --- | --- | --- |
| Organism | human | human | human |
| Data type | RNA-seq | snRNA-seq | RNA-seq |
| Samples | microdissected FFPE tumors | fresh frozen surgical resections | microdissected FFPE tumors |
| Comparison | MBM vs MNBM | MBM vs MNBM | MBM vs non tumor-bearing brain controls |
| Results availability | all genes | all genes | significant genes |
| Upregulated genes | 418 | 7201 | 685 |
| Downregulated genes | 568 | 9377 | 431 |
| Reference | PMID: 30787016 | PMID: 35803246 | PMID: 36435874 |

A large number of studies were identified for each neurodegenerative disease: 10 datasets for AD, 7 for PD and 4 for MS. Then we applied a robust meta-analysis methodology to identify genes consistently differentially expressed across datasets, as follows. Systematic review, data normalization and exploratory analyses have already been published (PD in PMID:36414996, MS in PMID:37023829, and AD in doi: https://doi.org/10.1101/2023.09.05.556293), and can be accessed in the web tool. A summary of this systematic review is provided in Supplementary Figure S1. We first performed differential gene expression analysis for each study, comparing *cases* (e.g. AD, PD or MS samples) vs *controls*. The number of significantly differentially expressed genes in each of these studiesis described in Supplementary Table S1. Next, we grouped the datasets according to the neurodegenerative disease and the respective brain regions of the samples examined, to perform five meta-analyses: AD cortex (AD-CT), AD hippocampus (AD-HP), PD substantia nigra (PD-SN), PD striatum (PD-ST), and MS. These analyses represent the genes differentially expressed in each of these brain regions in normal vs diseased tissue. Table 2 summarizes the technical characteristics and the results of each meta-analysis.

**Table 2.**
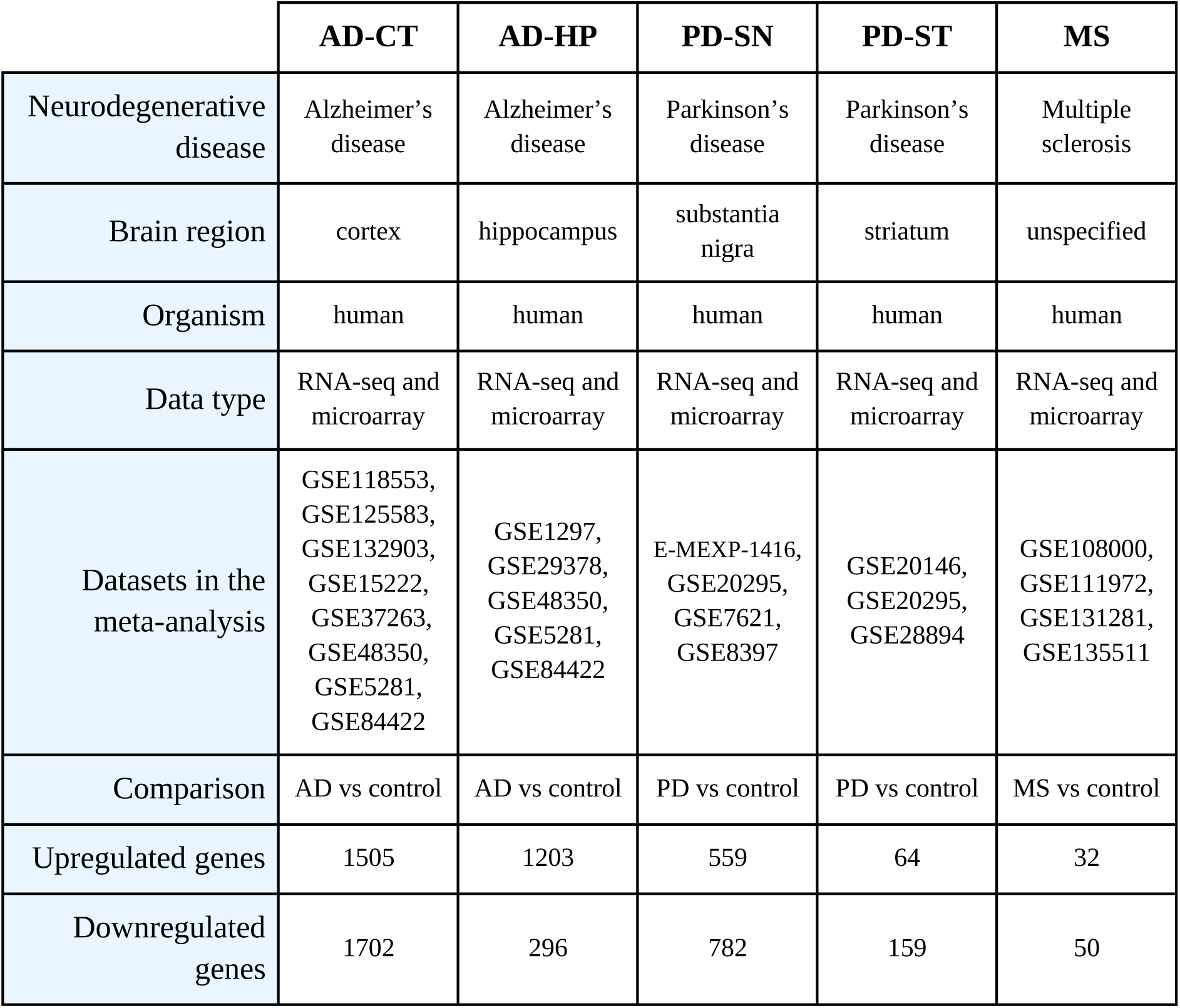
Results of meta-analysis of transcriptomics datasets of human samples of neurodegenerative diseases. Upregulated genes refer to the number of significant genes more expressed in the first term of the comparison (logFC > 0), and downregulated genes to those more expressed in the second term of the comparison (logFC < 0). AD: Alzheimer’s disease; CT: cortex; HP: hippocampus PD: Parkinson’s disease; SN: substantia nigra; ST: striatum; MS: multiple sclerosis.

|  | AD-CT | AD-HP | PD-SN | PD-ST | MS |
| --- | --- | --- | --- | --- | --- |
| Neurodegenerative disease | Alzheimer's disease | Alzheimer's disease | Parkinson's disease | Parkinson's disease | Multiple sclerosis |
| Brain region | cortex | hippocampus | substantia nigra | striatum | unspecified |
| Organism | human | human | human | human | human |
| Data type | RNA-seq and microarray | RNA-seq and microarray | RNA-seq and microarray | RNA-seq and microarray | RNA-seq and microarray |
| Datasets in the meta-analysis | GSE118553, GSE125583, GSE132903, GSE15222, GSE37263, GSE48350, GSE5281, GSE84422 | GSE1297, GSE29378, GSE48350, GSE5281, GSE84422 | E-MEXP-1416, GSE20295, GSE7621, GSE8397 | GSE20146, GSE20295, GSE28894 | GSE108000, GSE111972, GSE131281, GSE135511 |
| Comparison | AD vs control | AD vs control | PD vs control | PD vs control | MS vs control |
| Upregulated genes | 1505 | 1203 | 559 | 64 | 32 |
| Downregulated genes | 1702 | 296 | 782 | 159 | 50 |

### Melanoma brain metastasis-specific genes are often found dysregulated in multiple neurodegenerative diseases

Genes differentially expressed by metastatic cells in the brain compared to other metastatic locations that are also found dysregulated in neurodegenerative diseases may be indicative of shared cellular maladaptations to injury or inflammation within the CNS. Thus, we searched for differentially expressed genes in MBM-1 and MBM-2 (i.e., MBM vs MNBM) also found consistently altered in either neurological condition, as indicated by our meta-analyses (AD-CT, AD-HP, PD-SN, PD-ST and MS) (Supplementary Figure S2A). We identified genes with four possible behaviors (Figure 2A): consistent patterns, in which genes are upregulated (pattern 1) or downregulated (pattern 2) in both MBM and the corresponding neurodegenerative disease; and divergent patterns, where genes are upregulated in one disease and downregulated in the other (patterns 3 and 4). For both consistent and divergent patterns, we identified which genes are specific to one neurodegenerative disease and region (i.e., significant in a single meta-analysis), and which are found across several neurodegenerative diseases (Supplementary Figure S2B).

**Figure 2.**
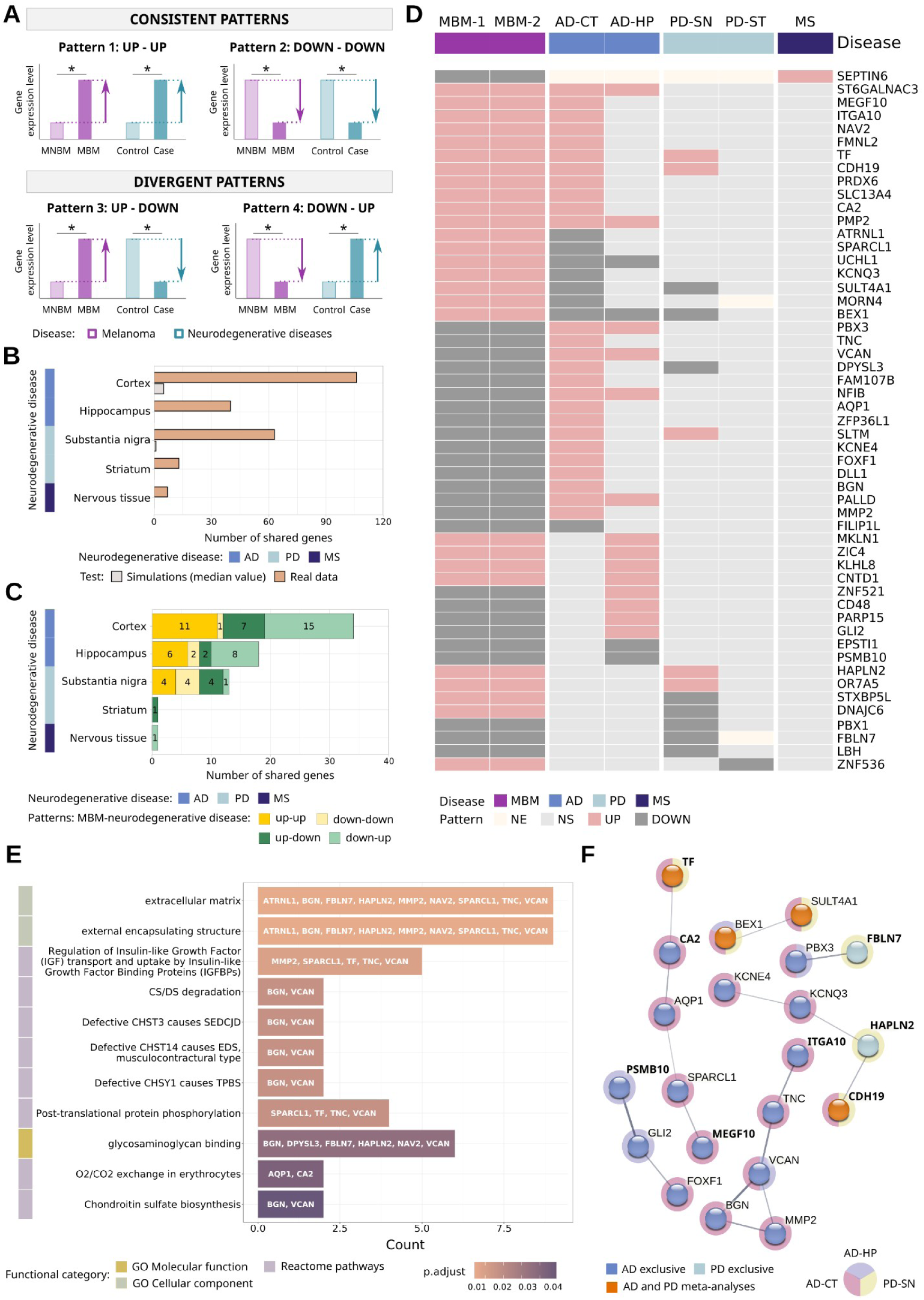
Neurodegenerative signature of melanoma brain-specific metastasis. (A) Qualitative representation of the patterns exhibited by significant genes. (B) Number of significant genes resulting from resampling analysis. (C) Distribution and (D) gene dysregulation profile by study/meta-analysis assessed. (E) Functional categories, whereX-axis represents the number of genes found in each category. (F) Protein-protein interaction network. Nodes represent genes, colored according to the neurodegenerative disease where it is significant. Normal text: consistent patterns; bold text: divergent patterns. Edge thickness indicates the structural and functional confidence of the interaction. Non-connected nodes are not shown. MBM: melanoma brain metastasis; MNBM: melanoma non-brain metastasis. AD: Alzheimer’s disease; CT: cortex; HP: hippocampus PD: Parkinson’s disease; SN: substantia nigra; ST: striatum; MS: multiple sclerosis; NE: not evaluated; NS: not significant. GO: gene ontology.

To assess robustness of the detected signal, we selected the significant genes shared between MBM-1 and MBM-2, hereafter referred to as MBM-1-2 genes. We looked for MBM-1-2 genes that were also dysregulated in neurodegenerative diseases. Resampling analysis, where we simulate the intersections with all the evaluated genes, showed that for each of the intersections between MBM-1-2 and neurodegenerative disease the number of significant common genes detected is higher than expected by random chance (Figure 2B), suggesting a shared biological signal between the two disease types. We next delved into the distribution of MBM-1-2 genes by neurodegenerative disease and pattern (Figure 2C). By merging these results we obtained a ‘MBM-1-2 neurodegenerative’ signature, composed by a panel of 53 genes (Figure 2D), enriched in 11 functional terms (Figure 2E). Of them, those associated with molecular functions and cellular components point to alterations of the extracellular matrix. The remaining terms are Reactome pathways mostly driven by VCAN and BGN genes, encoding two proteoglycans, versican and biglycan respecting. We also generated the protein-protein interaction (PPI) network (Figure 2F), which has significantly more interactions than expected (PPI enrichment p-value: 0.00111). Among the connected proteins in the network 19 are dysregulated in AD, 6 in PD and none in MS. In addition, genes with consistent and divergent patterns are connected, without generating specific clusters. Functional characterization of genes with consistent and divergent patterns separately, as well as genes by neurodegenerative disease can be found in the web tool.

We then explored the functional profile of the signature by analyzing concurrently the patterns and neurodegenerative diseases (e.g. genes with Pattern 4 between MBM-1-2 and AD-CT, being the cortex genes down-up in Figure 2C). AD Pattern 1, which consists of upregulated genes in both MBM-1-2 and the AD-CT, identified genes involved in cell adhesion molecule binding; (i.e., CDH19, FMNL2, ITGA10 and PRDX6). It is noteworthy that genes with Pattern 1 in the hippocampal region ST6GALNAC3 related to glycosylation and PMP2 to lipid metabolism) are also upregulated in AD-CT. AD Pattern 4 encompasses the major functional implications, with downregulated genes in MBM-1-2 and upregulated genes in AD (Figure 3). For both AD-CT and AD-HP regions genes are mostly involved in neurogenesis, development and the extracellular matrix; but in different proportions. Within the genes implicated in neurogenesis, we mainly find those specific to the hippocampus (ZNF521 and GLI2), and those common to both brain regions (VCAN, PBX3, PALLD and NFIB). In contrast, among development enlisted genes we identify a cortex-specific core, with four of them only associated with development functions (ZFP36L1, KCNE4, FOXF1 and AQP1). The term matrix comprises fewer genes, being retrieved from both cortex and hippocampus.

**Figure 3.**
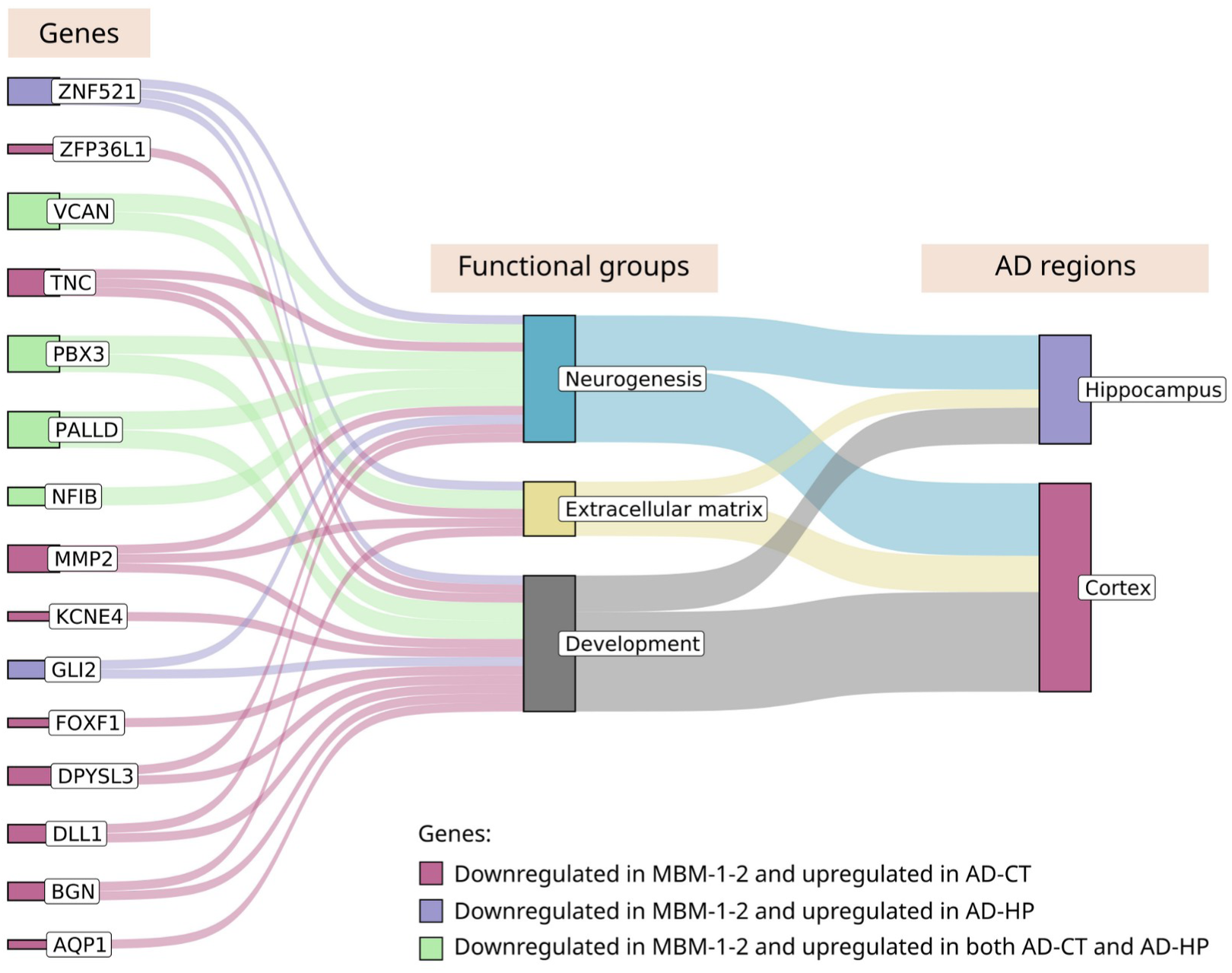
Functional classification of genes from MBM-1-2 and AD meta-analyses belonging to pattern 4. Functional groups attributed to the genes downregulated in MBM-1-2 and upregulated in AD-CT and/or AD-HP. Three categories are established (genes, functions, and brain regions). The size of the bars is proportional to the total number of genes found in each feature. AD: Alzheimer’s disease; CT: cortex; HP: hippocampus.

The functional significance of MBM-1-2 metastatic genes shared with PD are those related to the substantia nigra. Among upregulated genes in both diseases (Pattern 1), CDH19, HAPLN2, and OR7A5 are associated with adhesion and signaling transduction. Glycosaminoglycan binding genes (DPYSL3 and FBLN7) are present in Pattern 2, downregulated in both diseases. Of note, all genes account on Pattern 3, which are upregulated in MBM-1-2 and downregulated in PD-SN, have neuronal-specific features: BEX1 (potential glioblastoma biomarker), STXBP5L (neurotransmitter release), DNAJC6 (neuronal endocytosis) and SULT4A1 (neurotransmitter metabolism).

### Common genetic features between melanoma brain metastasis tumoral signature and neurodegenerative diseases profiles

Next, we identified which genes of the MBM tumor profile are also dysregulated in neurodegenerative diseases. We compared the differential gene expression results obtained in MBM-3 (*MBM vs brain controls* comparison) with the results of the five meta-analyses (AD-CT, AD-HP, PD-SN, PD-ST and MS). In total, 195 significant genes were spotted finding all four possible patterns (Figure 4A). The highest number of shared genes are reported in the intersections of MBM-3 with AD-CT, AD-HP and PD-SN (Figure 4B). Regarding the neurodegenerative disease, 112 were AD-specific, 57 PD-specific, 7 MS-specific and 19 were dysregulated in both AD and PD (Figure 4C). Among the total genes, those belonging to consistent patterns stand out. Within the significant PPI network of Pattern 1 (PPI enrichment p-value: 2.79e-08), we encountered the following clusters (Figure 4D Pattern 1): translation related genes (RPL13A, NACA, RPL35A, RPS9, RPS17, GSUB, PABPC1 and EIF4B), mostly associated with PD; development and cell differentiation genes (MAP3K1, F2R, RHOA, CAV1, MSN, ITGB3, S100A4, ANXA2, TGFBI, GYPC, COL3A1, AEBP1, CRTAP, MGP, PLOD3, MLEC, CHST14, CSPG4, LAMB2, ITGA10), mostly associated with AD; and chromatin remodeling and nucleosome organization genes (HIST1H2AB, HIST1H3D, HIST1H2BK, HIST1H2AD, HIST1H4L, ACTL6A, ZNF217, HDAC1), with similar proportions associated with AD and PD. Genes constituting Pattern 2 are predominantly dysregulated in AD. They also generate a significant network (PPI enrichment p-value: 0.00015) (Figure 4D Pattern 2), composed by genes linked with the synaptic process, particularly those contributing to vesicle formation. The PPI networks of convergent Pattern 3 and Pattern 4, as well as the functional characterisation of the genes by neurodegenerative disease, can be explored in the web tool.

**Figure 4.**
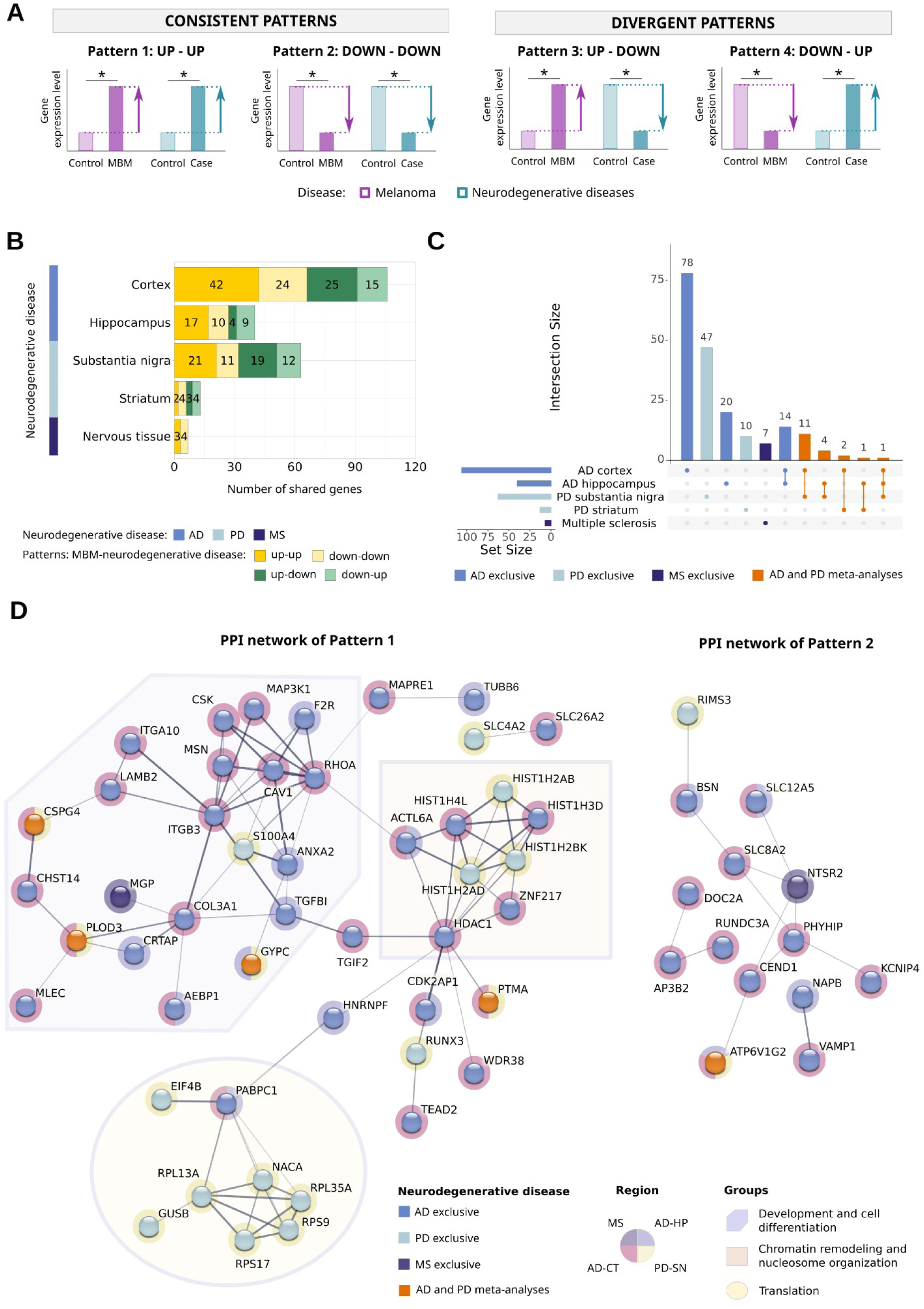
Neurodegenerative signature for MBM tumoral profile. (A) Qualitative representation of the patterns exhibited by significant genes in MBM-3 and neurodegenerative diseases meta-analyses. (B) Number of significant genes of the signature separated by neurodegenerative disease and pattern. (C) Upset plot of the signature. Representation of the total number of significant genes by neurodegenerative disease (horizontal bars), and the fraction present in the intersection of the designated groups (vertical bars), identified by colored dots beneath. (D) Protein-protein networks for consistent genes between MBM-3 and neurodegenerative diseases: Pattern 1 (upregulated genes), Pattern 2 (downregulated genes). Nodes correspond to genes, with color-coded based on their significance in the neurodegenerative diseases. Edge thickness indicates the structural and functional confidence of the interaction. Non-connected nodes have been discarded.

### DNAJC6 and ITGA10 as potential key genes in the neurobiology of melanoma brain metastases

To identify genes that were not only differentially expressed in MBM compared to control tissue (MBM-3 genes) but also specific to brain metastasis (MBM-1-2 genes), we performed an intersection analysis to identify significant genes in all three MBM differential gene expression results. We then determined which of these genes were also dysregulated in neurodegenerative diseases. We found two genes that meet these gene expression patterns: ITGA10, which is upregulated in MBM-1, MBM-2, MBM-3 and AD-CT; and DNAJC6, which is upregulated in MBM-1 and MBM-2, whilst is downregulated in MBM-3 and PD-SN.

### Web tool

The metafun-mbm web resource (https://bioinfo.cipf.es/metafun-mbm/) includes the detailed results of each section developed in this manuscript, offering free access to users for confirmation of the described findings. The interactive interface allows users to explore the intermediate results generated to obtain the dysregulated profiles of MBM and the neurodegenerative diseases (i.e. systematic review, exploratory analysis, differential expression and meta-analysis). Users will also be able to explore the neurodegenerative signature of MBM-1-2 and MBM-3; together with the functional profiles at the different levels presented in this manuscript.

## DISCUSSION (Results and Discussion may be combined into one section)

Cancer neuroscience is increasingly appreciated as a multidisciplinary avenue of discovery for both contributing fields (PMID: 32302564; PMID: 37059069). Studies rooted in brain metastasis biology have identified subpopulations of glial cells and revealed their capabilities (PMID: 27225120; PMID: 29892069; PMID: 33953400). Indeed, ongoing clinical trials combine neuroactive drugs with anticancer standards of care to exploit the relationship between neuronal activity and tumor cells (PMID: 35493943). With our study we aim to highlight potential such lines of investigation for both melanoma brain metastasis and neurodegeneration by explicitly identifying common molecular mechanisms.

Thus, we conducted an *in silico* analysis of public transcriptomic data, retrieved from screening scientific literature following the FAIR principles (Findable, Accessible, Interoperable, Reusable) (https://doi.org/10.1038/sdata.2016.18). This approach provided us with the state of the art of transcriptomic studies of MBM and neurodegenerative diseases. We have implemented meta-analysis methodology when possible, integrating data from different datasets to increase the robustness of the analysis. However, we cannot incorporate all the identified studies from the literature. The lack of homogeneity in both the technical and clinical variables, along with differences in the phenotype and treatments of the samples, limits the inclusivity of studies within the meta-analysis. Sufficient data have been obtained to perform AD-CT, AD-HP, PD-SN, PD-ST and MS meta-analyses. For MBM, which is characterized by a scarcity of research, we have not achieved the minimum number of necessary studies. Therefore, we have characterized the transcriptomic profiles from individual analyses.

An important factor to consider in our analysis is the purposeful “apples-to-oranges” approach: we have compared changes in *tumors* composed of melanoma cells to changes in *brain tissue* in neurodegenerative pathology. This strategy was taken in order to further explore brain metastatic melanoma cells’ capacity for neuro-mimicry, identified by multiple works in the field (PMID: 35262173; PMID: 35803246). Despite the inherent differences between the tissues analyzed, common molecular changes were observed, discussed below. Nonetheless future work may center on comparisons between the altered *brain microenvironments* in MBM and neurodegenerative diseases. Particularly useful may be comparisons of single cell-level data, as the brain is a complicated ecosystem of cell types -- each with a role to play in disease progression -- and we have made comparisons using bulk RNA-sequencing of brains with AD/PD. Inappropriate neuroinflammation is thought to drive neurodegenerative pathology, and melanoma cells have developed mechanisms to quell neuroinflammation to survive in the BME. By examining this latter dynamic through the lens of both melanoma cell gene expression (as in this work) and that of immune cells and glia, light may be shed on the biology of neurodegeneration as well. Due to a paucity of samples worldwide, our MBM data are agnostic to brain geography whereas our AD and PD data are coarsely defined by brain region. Of course the brain has extensive spatial heterogeneity of gene expression, and this would be an even more important factor to consider in comparing the MBM-altered brain to AD and PD-bearing brains.

Another important limitation of this work is the inclusion only of MBM data. Non-small cell lung cancer and breast cancer give rise to brain metastasis in greater numbers and may also exhibit molecular commonalities with neurodegeneration. Indeed, breast cancer brain metastatic cells can engage in synapses with neurons, suggesting a degree of neuro-mimicry (PMID: 31534217). We postulate that melanoma may have more in common with neurodegeneration than other cancers due to its developmental history in the neural crest, but this cannot be proven without a thorough investigation of other sources of brain metastasis.

MBM gene expression profiling compared to non-brain metastases revealed a significant enrichment for common genes with neurodegenerative pathologies (Figure 2). Despite the established epidemiologic connection between melanoma and Parkinson’s disease, MBM signatures had more in common with those of AD. *DPYSL3* is of note because it is downregulated in MBM, upregulated in AD-CT, and downregulated in PD-SN (Figure 2). DPYSL3 was identified by the Clinical Proteomic Tumor Analysis Consortium as a specific marker of claudin-low triple-negative breast cancers, and DPYSL3 knockdown slowed proliferation in breast cancer cell lines while increasing motility and markers of the epithelial-to-mesenchymal transition (PMID: 30498031). Dpysl3 knockdown in rat microglia inhibited their migration, activation, and phagocytic activity (PMID: 23988434). The (PMID: 23193282) of human single cell RNA-sequencing data shows that DPYSL3 may be expressed by a few neuronal subtypes as well as astrocytes (data not shown). Future work may reveal a role for DPYSL3 in melanoma, and could perhaps help to delineate mechanisms of microglial activation that are divergently regulated in AD and PD.

Throughout our analyses, extracellular matrix (ECM) regulation recurred as a point of congruence between neurodegenerative diseases and MBM (Figure 2E and Figure 3). As effort has been expended to characterize the cellular components of the BME, the ECM is an underexplored avenue in both fields despite constituting about 20% of brain volume (PMID: 36744003). The common constituents of the ECM dysregulated across diseases include BGN, or biglycan (Figure 2). Bgn knockout in mice causes decreased memory performance and brain metabolic dysfunction, and *BGN* is upregulated specifically in dormant cancer cells in the brain from a breast cancer brain metastasis *in vivo* model (PMID: 37456245). The secreted glycoprotein SPARCL1 was upregulated in 2 MBM cohorts and decreased in expression in AD. SPARCL1 also exhibited increased expression in dormant breast cancer brain metastasis cells compared to proliferating cells (PMID: 37456245) and has been implicated in progression of various cancers (PMID: 28010899). Investigation into the role of these and other ECM components commonly dysregulated in the diseased BME may start to unlock the role of the ECM overall in these pathologies. Monitoring of CSF for proteolytic cleavage products of brain-derived ECM molecules has been suggested as a monitoring strategy for conditions such as ALS and epilepsy; a similar approach could be undertaken for the diseases analyzed here if the ECM indeed proves critical to progression (PMID: 35480959).

Transferrin, or TF, was also upregulated across AD cortex samples, PD substantia nigra samples, and 2 MBM cohorts. Iron homeostasis is critical to normal brain functioning, but excess iron can precipitate oxidative damage. TfR1-mediated Tf-Fe endocytosis at the blood-brain barrier is a decisive bottleneck in iron uptake. Increased iron levels have been observed in AD, PD, and Huntington’s disease, but whether this is a cause or a consequence of disease progression is not clear (https://doi.org/10.1111/brv.12521).

The findings from this research contribute to the characterisation of the MBM neuronal patterns. The unveiled dysregulated genes, together with their associated biological functions and signaling pathways, represent potential biomarkers to further advance in the profiling of MBM and in the understanding of clinical outcomes.

## MATERIALS & METHODS

The bioinformatics analysis was conducted with version 4.1.2. of the R programming language (R Core Team (2021). R: A language and environment for statistical computing. R Foundation for Statistical Computing, Vienna, Austria. URL https://www.R-project.org/.). The implemented packages and their corresponding versions can be consulted in the Supplementary Table S2.

### Data collection

Literature screening was conducted during December 2022. Regarding the disease addressed, all selected studies met the following inclusion criteria: i) type of data: transcriptomic data, ii) organism: human (*Homo sapiens*), iii) type of biological sample processing: frozen *postmortem* tissue sections. Additional specifications differed for each type of disease. MBM data corresponded to differential gene expression results from MBM versus melanoma non-brain metastasis. To broaden the analysis, MBM versus non tumor-bearing brain controls results were also collected. For neurodegenerative diseases, the systematic review has already been published in (MS in PMID: 37023829, and PD in PMID: 36414996).

### Differential gene expression and meta-analyses in neurodegenerative diseases studies

Novel differential gene expression analysis was performed independently for each selected study of each neurodegenerative disease. Case versus control comparison was tested by implementing the linear regression model of the R limma package (PMID: 25605792). When necessary, batch effect was included as covariable. P-values were adjusted by the Benjamini and Hochberg (BH) procedure (PMID: 11682119), considering a significant change in gene expression when FDR (False Discovery Rate) < 0.05. Log2 Fold Change (logFC) was calculated to define the direction and magnitude of change. Thus, logFC positive genes are upregulated in cases (downregulated in controls), and logFC negative genes are upregulated in controls (downregulated in cases).

Then results were integrated into five meta-analyses based on the neurodegenerative disease and the brain region examined (AD-CT, AD-HP, PD-SN, PD-ST and MS). Meta-analyses were developed as previously described (García-García F. Methods of functional enrichment analysis in genomic studies [PhD Thesis]. 2016). To account for the individual study heterogeneity, we implemented DerSimonian and Laird random-effects model (PMID: 3802833) using metafor R package (https://doi.org/10.18637/jss.v036.i03). Meta-analyses statistics were calculated, and p-values were corrected by BH method (PMID: 11682119). FDR’s significant cutoff was established as 0.05.

### Intersection analysis

Significant genes for MBM differential gene expression results and neurodegenerative diseases meta-analyses were selected by setting an FDR of 0.05, and classified based on the direction of change in the corresponding statistical comparison: upregulated genes (logFC > 0) and downregulated genes (logFC < 0). We then looked for common significant genes between MBM and each neurodegenerative disease by performing intersection analyses of the resulting gene lists. Due to the availability of two studies with the MBM vs melanoma non-brain metastasis comparison, we filtered out genes with the same pattern in MBM-1 and MBM-2 studies. To visualize the results, we elaborate bar plots with the R package ggplot2 (Wickham H (2016). ggplot2: Elegant Graphics for Data Analysis. Springer-Verlag New York. ISBN 978-3-319-24277-4, https://ggplot2.tidyverse.org), upset plots with the R package UpSetR (PMID: 28645171), and heatmaps with the R package ComplexHeatmap (https://doi.org/10.1002/imt2.43).

### Resampling

We performed a resampling analysis to detect whether the genes resulting from the intersection analysis had non-arbitrary biological signals or had arisen randomly. For each pair of MBM and neurodegenerative disease results we first selected the common genes assessed in both diseases. Then we performed 10000 iterations of the following process: (1) arbitrarily select the number of genes identified in the intersection analysis and (2) identify how many of these genes are significant in the two MBM studies. Finally, we calculated the median of the calculated simulations.

### Functional signatures of the common transcriptomic features

The unveiled gene profiles were fully functionally characterized by two strategies: protein-protein interaction analysis (PPI) and overrepresentation analysis (ORA). PPI analyses were conducted using the STRING R package (PMID: 30476243). The interaction networks were generated using default parameters, where we evaluated both functional and physical protein associations. Significant networks were considered when PPI enrichment p-value < 0.05. For the purpose of visualization, disconnected proteins were hidden, and their interaction confidence value represented the strength of the interactions.

We also screened the biological relationships in the selected gene sets. Biological annotations were obtained using the AnnotationDbi R package (Pagès H, Carlson M, Falcon S, Li N (2023). AnnotationDbi: Manipulation of SQLite-based annotations in Bioconductor. R package version 1.62.1) from the org.Hs.eg.db R package (Carlson M (2019). org.Hs.eg.db: Genome wide annotation for Human. R package version 3.8.2.): i) Reactome pathways, ii) KEGG pathways, iii) GO Biological Processes, iv) GO Molecular Functions, and v) GO Cellular Components; being the last three from the Gene Ontology database. To perform the functional tests, the statistical method ORA was implemented with the clusterProfiler R package (PMID: 34557778). We analyzed those gene sets within the range of 10 (minimal size) and 500 (maximal size) genes, thus filtering out specific and general results. We calculated GeneRatio, BgRatio, q-value and p-value statistics. P-values were adjusted by BH method (PMID: 11682119), considering statistical significance when FDR < 0.05. Plots were generated with the corresponding functions of the enrichplot R package (Yu G (2023). enrichplot: Visualization of Functional Enrichment Result. R package version 1.20.0).

### Web tool

The metafun-mbm web tool (https://bioinfo.cipf.es/metafun-mbm/) is an open resource to explore in-depth the data and results presented in this manuscript. It has been developed with Quarto system (https://github.com/quarto-dev/quarto-cli), providing an user-friendly and interactive environment to navigate through the web. Plots have been generated with ggplot2 (Wickham H (2016). ggplot2: Elegant Graphics for Data Analysis. Springer-Verlag New York. ISBN 978-3-319-24277-4, https://ggplot2.tidyverse.org), plotly (Sievert C (2020). *Interactive Web-Based Data Visualization with R, plotly, and shiny*. Chapman and Hall/CRC. ISBN 9781138331457, https://plotly-r.com) and enrichplot (Yu G (2023). enrichplot: Visualization of Functional Enrichment Result. R package version 1.20.0) R packages. Results are displayed in seven sections: i) overview of the implemented pipeline, ii) MBM expression profiles, iii) neurodegenerative diseases expression profiles, iv) neurodegenerative signature and functional profiling of brain metastasis-specific signatures (MBM-1-2 results), v) neurodegenerative signature and functional profiling of MBM tumor profile (MBM-3 results), and vi) methods description.

## ACKNOWLEDGEMENTS

The authors thank the Principe Felipe Research Center (CIPF) for providing access to the cluster, co-funded by European Regional Development Funds (FEDER) to the Valencian Community 2014-2020.

This research was supported by and partially funded by the Institute of Health Carlos III (project IMPaCT-Data, exp. IMP/00019), co-funded by the European Union, European Regional Development Fund (ERDF, “A way to make Europe”), PID2021-124430OA-I00 funded by MCIN/AEI/10.13039/501100011033/ FEDER, UE (“A way to make Europe”). Irene Soler-Sáez is supported by a predoctoral grant FPU20/03544 funded by the Spanish Ministry of Universities. We also thank the authors of the previous publications (PMID: 30787016; PMID: 35803246; PMID: 36435874) for their commitment to open science by depositing the data in public repositories, making the data available and reusable.

## AUTHOR CONTRIBUTIONS

Conceptualization: EH, FGG; Data Curation: AK, ISS, ALC, JFCS; Investigation: EH, FGG, AK, ISS, MIV; Bioinformatic Analysis: ISS, ALC, BGC, JFCS; Supervision: EH, FGG, MRH; Writing-Original Draft Preparation: EH, FGG, AK, ISS, MIV, BGC, ALC, JFCS; all authors read and approved the final manuscript.

---

**Figure S1.**
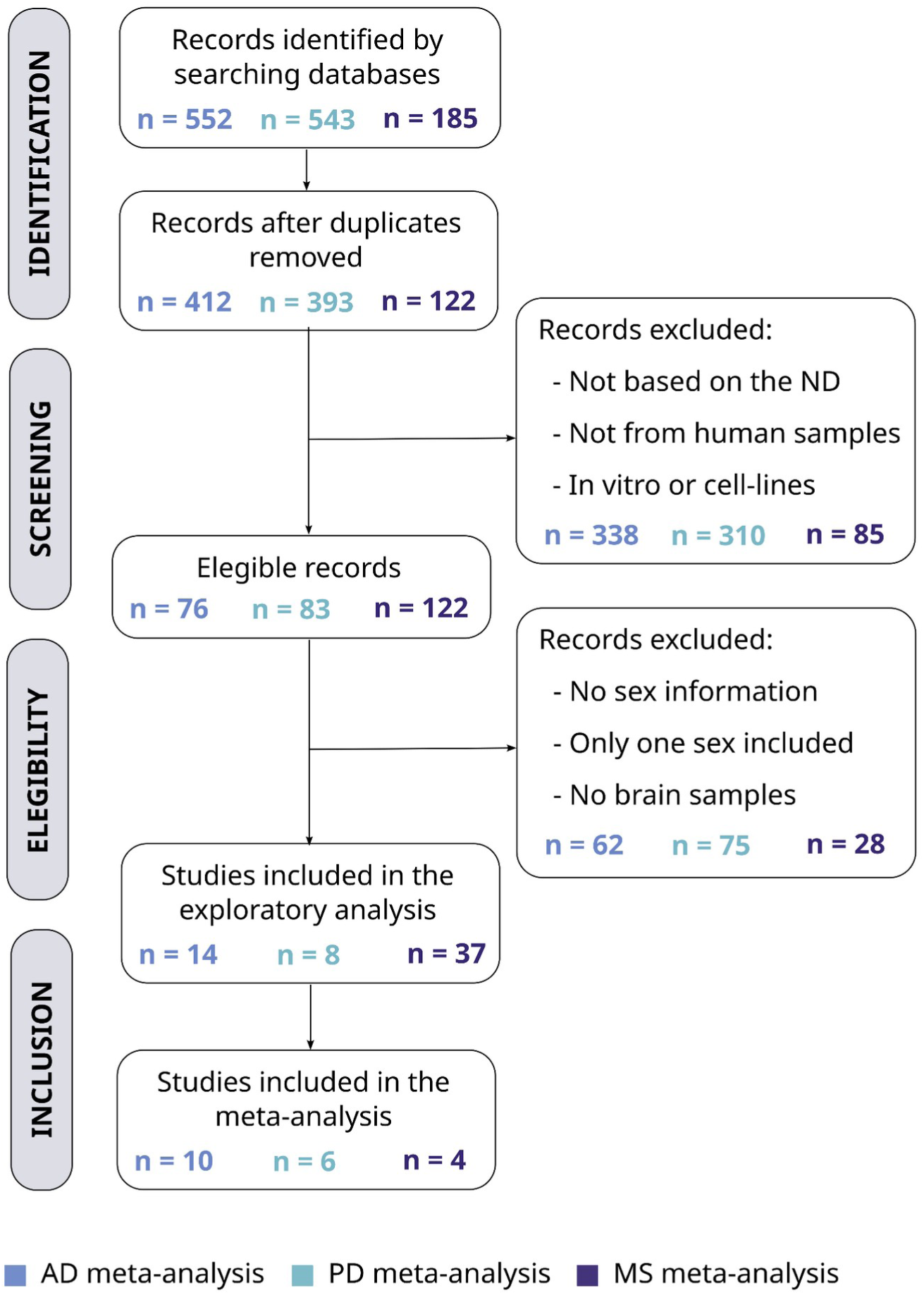
Diagram of the systematic reviews conducted for each neurodegenerative disease: Alzheimer’s disease, Parkinson’s disease and multiple sclerosis. Multistep systematic review according to PRISMA statement guidelines (PMID: 19621072). The number of retained or discarded studies are indicated at each phase (n) for each neurodegenerative disease (defined by color). PRISMA: Preferred Reporting Items for Systematic Reviews and Meta-Analyses; ND: neurodegenerative disease; AD: Alzheimer’s disease; PD: Parkinson’s disease; MS: multiple sclerosis.

**Figure S2.**
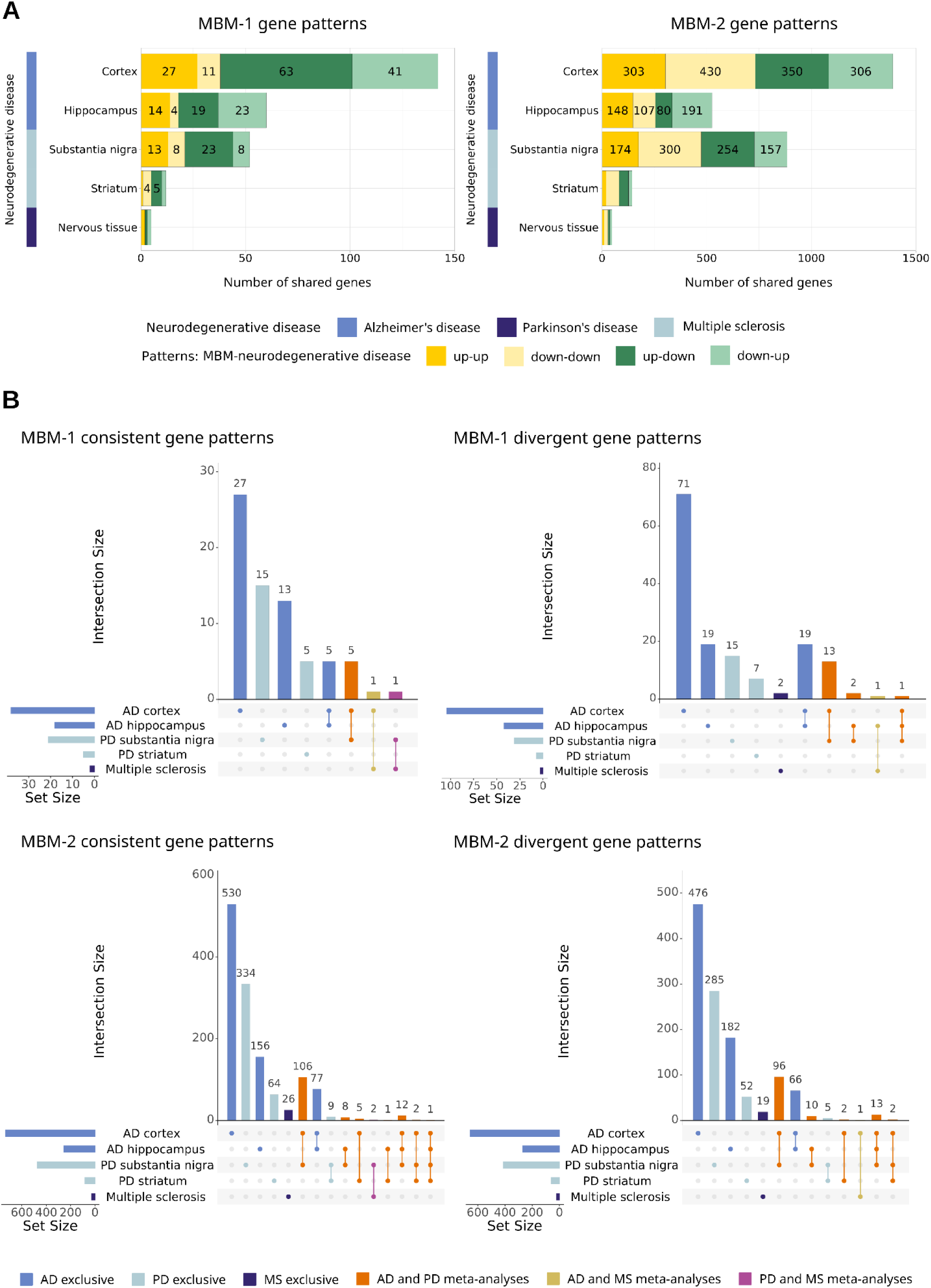
Individual intersection analyses for MBM-1 and MBM-2. (A) Distribution of the number of dysregulated genes in both MBM and each neurodegenerative disease categorized by pattern (MBM-1 right panel, MBM-2 left panel). (B) Genes from A) were classified based on their significance in the neurodegenerative disease meta-analyses and whether they display consistent or divergent patterns. MBM: melanoma brain metastasis; AD: Alzheimer’s disease; PD: Parkinson’s disease; MS: multiple sclerosis.

**Table S1.** Number of significantly differentially expressed genes in diseased vs normal tissue per study. The study identifier of the public repository, the neurodegenerative disease, and the brain region assessed are indicated. The number of differentially expressed genes in cases vs. controls is indicated, with upregulated genes being those more expressed in cases (logFC > 0), and downregulated genes those more expressed in controls (logFC < 0). ID: identifier; AD: Alzheimer’s disease; CT: cortex; HP: hippocampus PD: Parkinson’s disease; SN: substantia nigra; ST: striatum; MS: multiple sclerosis.

| Study ID | Neurodegenerative disease | Brain region | Upregulated genes | Downregulated genes |
| --- | --- | --- | --- | --- |
| GSE118553 | AD | CT | 1858 | 1367 |
| GSE125583 | AD | CT | 5706 | 5303 |
| GSE132903 | AD | CT | 7631 | 6627 |
| GSE15222 | AD | CT | 4859 | 3756 |
| GSE37263 | AD | CT | 0 | 0 |
| GSE48350 | AD | CT | 61 | 64 |
| GSE5281 | AD | CT | 1200 | 5012 |
| GSE84422 | AD | CT | 439 | 455 |
| GSE1297 | AD | HP | 1 | 0 |
| GSE29378 | AD | HP | 801 | 497 |
| GSE48350 | AD | HP | 6092 | 1856 |
| GSE5281 | AD | HP | 1132 | 902 |
| GSE84422 | AD | HP | 0 | 0 |
| E-MEXP-1416 | PD | SN | 0 | 0 |
| GSE20295 | PD | SN | 42 | 32 |
| GSE7621 | PD | SN | 0 | 2 |
| GSE8397 | PD | SN | 2001 | 1381 |
| GSE20295 | PD | ST | 0 | 0 |
| GSE20146 | PD | ST | 0 | 0 |
| GSE28894 | PD | ST | 90 | 144 |
| GSE108000 | MS | undefined | 2038 | 3114 |
| GSE111972 | MS | undefined | 0 | 0 |
| GSE131281 | MS | undefined | 162 | 188 |
| GSE135511 | MS | undefined | 451 | 401 |

**Table S2.** List of R packages and versions implemented in the bioinformatic workflow.

| Package | Version |
| --- | --- |
| ggplot2 | 3.4.0 |
| UpSetR | 1.4.0 |
| stringr | 1.4.0 |
| ggpubr | 0.4.0 |
| ggh4x | 0.2.3 |
| tidyverse | 1.3.1 |
| dplyr | 1.0.8 |
| tidyr | 1.2.0 |
| clusterProfiler | 4.2.2 |
| org.Hs.eg.db | 3.14.0 |
| AnnotationDbi | 1.56.2 |
| egg | 0.4.5 |
| reshape2 | 1.4.4 |
| ComplexHeatmap | 2.13.4 |
| enrichplot | 1.14.2 |

